# Septins disruption controls tumor growth and enhances efficacy of Herceptin

**DOI:** 10.1101/2020.02.19.954529

**Authors:** Rakesh K Singh, Kyu Kwang Kim, Negar Khazan, Rachael B. Rowswell-Turner, Christian Laggner, Aaron Jones, Priyanka Srivastava, Virginia Hovanesian, Liz Lamere, Thomas Conley, Ravina Pandita, Cameron Baker, Jason R Myers, Elizabeth Pritchett, Awada Ahmad, Luis Ruffolo, Katherine Jackson, Scott A. Gerber, John Ashton, Michael T. Milano, David Linehan, Richard G Moore

## Abstract

Septin expressions are altered in cancer cells and exhibit poor prognoses in malignancies. As the first approach to develop a septin filament targeting agent, we optimized the structure of Forchlorfenuron (FCF), a known plant cytokinin to generate UR214-9, which contrary to FCF, causes septin-2/9 filamental structural catastrophe in cancer cells without altering cellular septin protein levels. *In-silico* docking using septin-2/septin-2 dimer complex showed that UR214-9 displaced the guanine carbonyl oxygen from the GDP binding domain and showed increased binding energy than FCF(−8.59vs-7.21). UR214-9 reduced cancer cell growth, downregulated HER2/STAT-3 axis and controlled growth of HER2+ pancreatic, breast and ovarian cancer xenografts in NSG mice and enhanced response of Herceptin against HER2+breast cancer xenograft. Transcriptome analysis of UR214-9 exposed cells demonstrated significant perturbation of <20 genes compared to afatinib which impacted >1200 genes in JIMT-1 breast cancer cells indicating target specificity and non-transcriptional functions of UR214-9. In summary, disrupting septins via UR214-9 is a new approach to control the growth of HER2+ malignancies.

## Introduction

Septins are a family of GTP-binding cytoskeletal proteins that participate in cytokinesis, cell migration, chromosomal dynamics and protein secretion. Septins hetero-oligomerize to generate scaffolding filaments, bundles, and rings within cells^1–11^. Additionally, septins are a critical cytoskeletal component that regulate the function of tubulin and actin. Altered septin protein expression in pancreatic, kidney, lung, colorectal, skin, brain, endometrial, ovarian, breast and other malignancies have been observed^12–16^. Aberrant septin expression has also been linked to neurodegenerative/neuromuscular diseases, blood disorders, infertility, and developmental disabilities^17–19^. It is unclear whether aberrant enrichment of individual septin family members is enough to enhance tumorigenesis or if a specific hetero-oligomer assembly may be implicated. Pharmacologic agents to target septins have remained elusive, largely because the oligomeric structural configurations of septins pose difficult challenges in designing therapies.

In this study, we investigated the impact of individual septins on the survival of patients with cancers of the pancreas, breast, lung, kidney, or liver cancer or with melanoma. To determine the effect of septins on survival, we used the Human Protein Atlas (HPA), and publically available transcriptional data and tools available at R2:Genomics Analysis and Visualization Platform^20^ (https://hgserver1.amc.nl/cgi-bin/r2/main.cgi). We describe a potent septin modulator, UR214-9, which disrupts structural organization of septin-2 and septin-9 as well as of β-actin, and controls cancer cell proliferation and tumor growth. We have employed molecular docking techniques to investigate how UR214-9 and its analogs interact with the elements of the GDP binding domain, and of the known FCF binding pocket. To identify how gene expression is impacted by UR214-9, and thereby characterize its off-target liabilities, we have conducted transcriptome analyses of UR214-9 treated breast and pancreatic cancer cells. In summary, our studies present UR214-9, as a potent and novel septin filamental modulator and demonstrate that the dismantling of septin structures in pancreatic, ovarian and breast cancer cells by UR214-9 can be an effective therapeutic strategy.

## Results

### Enrichment of septins correlates with decreased survival in patients with cancer

Publically accessible microarray data bases of pancreatic cancer and ovarian cancer patients deposited at R2:Genomics Analysis and Visualization Platform(https://hgserver1.amc.nl/cgi-bin/r2/main.cgi) were analyzed. Septin-2 mRNA was enriched in malignant pancreas compared to normal pancreas (Figure-1A, p=1.3e^−4^). Similarly, ovarian cancer epithelium expressed significantly enrichment of septin−2 compared to normal stroma (Figure-1B-left, p=1.2e^−7^). Micro-dissected stroma of malignant ovarian stroma was also exhibited elevated expression of septin-2 mRNA than normal stroma (Figure-1B-right, p=1.21e^−4^). Similarly, compared to normal stroma, tumor epithelium components of malignant breasts showed increased septin-2 mRNA enrichment (Figure-1C-left, p=0.49e^−3^). Further, increase in invasive area of breast tumors led to increased septin-2 enrichment (Figure-1C-right, p=8.9e^−3^). Kaplan-Meier survival of patients with pancreatic cancer, grouped by the extent of septin-2 expression (from microarray data available at https://hgserver1.amc.nl/cgi-bin/r2/main.cgi^20^ and Human Protein Atlas^21^, show that septin-2 mRNA enrichment significantly (p=0.0011) correlates with increased mortality (Figure 1D-left). Similarly, enrichment of septin-7 and −9 correlates with increased mortalities in pancreatic cancer patients (Supplementary Figure 1). Septin-2 enrichment is also an unfavorable factor for patients with breast (Figure-1D, middle, p=3.9e-3) and ovarian cancer (Figure-1D, right, p=0.011). Analysis of the survival prospects based on other septins indicate that septin-7 enrichment was found to be unfavorable for the patients diagnosed with malignancies of breast (*p*=0.0079) (Supplementary Figure-1).

**Figure-1:**
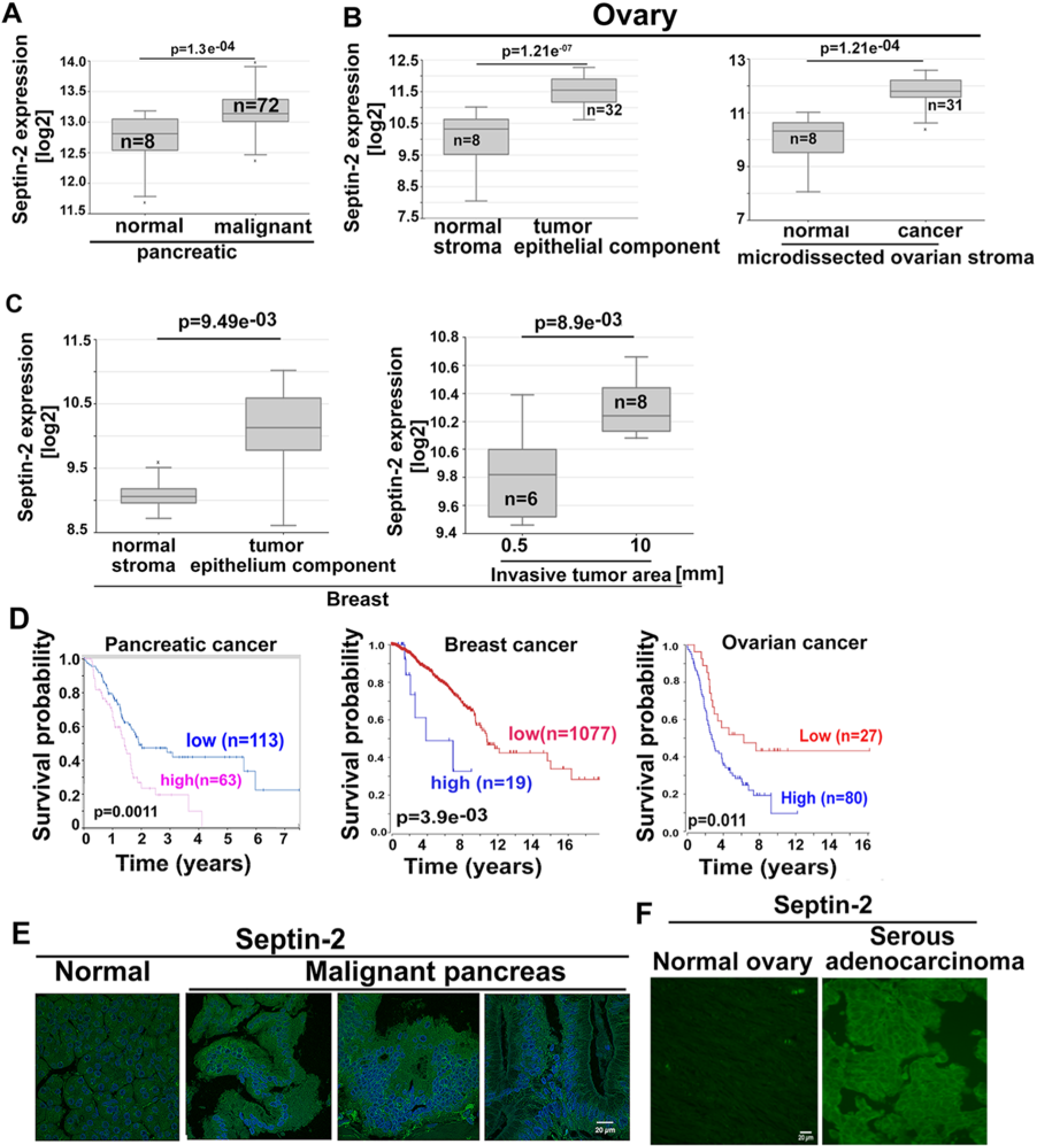
**(A-C)**: Septin-2 expression in normal and malignant pancreatic, ovarian and breast cancer tissues were analyzed using the publicly accessible patient’s tumor microarray data deposited on R2-Genomics Analysis and Visualization Platform (https://hgserver1.amc.nl/cgi-bin/r2/main.cgi). (**D**): Kaplan Meier survival analyses of the pancreatic, ovarian and breast cancer cancer patients using the data and tools available at the Human Protein Atlas or at R2-R2-Genomics Analysis and Visualization Platform show that septin-2 enrichment correlates with decreased survival among pancreatic, breast and ovarian cancer patients. Septin-7 and −9 enrichment was associated with increased mortality as well (see supplementary Figure-1). **(E):** Pancreatic Tumor microarray from US Biomax Inc (catalog number: T142a) was stained with septin-2 antibody (Sigma Aldrich Inc. catalog number: HPA018481), followed by sourced-matched secondary antibody (DyLight-488, catalog number DI-1488, Vector laboratories Inc.). Imaged were recorded as described in the methods section. Malignant tissues showed higher septin-2 expression than tissues isolated from normal pancreas. **(F):** Malignant serous ovarian cancer tissues showed increased septin-2 expression than normal ovaries. Ovarian tumor microarray from US Biomax Inc (catalog OV241a) was stained with septin-2 antibody (Sigma Aldrich catalog number: HPA018481) followed by a sourced-matched secondary antibody (DyLight-488, catalog number DI-1488, Vector laboratories Inc.). Imaged were acquired as described in method section.

### UR214-9 causes septin-2 catastrophe in cells

The chemical structure of UR214-9 is shown in Figure-2A. UR214-9 was obtained by the structure-activity relationship guided optimization of FCF. Incorporation of a group of fluorine atoms on phenyl ring and installation of a chlorine atom at C-6 of pyridine ring made UR214-9 a potent disruptor of septin’s filamental structure. Confocal microscopy at higher resolution (60×2) was employed to determine the impact of DMSO, FCF (+ve control) and UR214-9 on the structural arrangement of septin-2, 6, 7 and −9 in a panel of BXPC-3, CAPAN-1, Panc-1 (pancreatic) and JIMT-1(breast) and SKOV-3 ovarian cancer cells. While FCF seems to strengthen the septin-2 filaments in BXPC-3 cells (Figure-2B, left-lower), Septin-2 needles in PANC-1 were disarranged and translocated at the cell-surface after UR214-9 treatment (1uM) (Figure 2C, lower left). Similarly, the septin-2 needles in JIMT-1 cells after drug treatment showed structural disruptions and relocation to nuclear periphery (Figure-2C, right-lower). Next, the confocal microscopy was employed to investigate the response of other septin family members in PANC-1 cells upon treatment with UR214-9. Septin-7 showed reduced expression whereas septin-9 showed disarrangement of filamental structure (Supplementary Figure-2). Septin-4, −6, did not exhibit clear filamental structures and showed punctate staining instead, which was either reduced in the treatment group compared to DMSO treated control or the drug effect was inconclusive (data not shown). Similarly, UR214-9 treated JIMT-1 breast cancer cells showed strong structural disarrangement and relocation of septin-2 on the periphery of nucleus (Figure-2, right). JIMT-1 cells did not exhibit defined septin-7 structures, and therefore, the effect of UR214-9 on septin-7 remains ambiguous (Supplementary Figure-2, lower left). However, the confocal microscopy of PANC-1 and JIMT-1 cells treated with UR214-9 exhibited clear disarrangement in septin-9 filament structures (Supplementary Figure-2, lower right). Whether UR214-9 treatment alters expression of septin family of proteins was investigated by immunoblotting the total cell-lysates of PANC-1, MDA-MB-231, JIMT-1 and MCF-7 cancer cells. The immunoblots were probed with validated septin-2, 6,7 and −9 antibodies. In PANC-1 cells, septin-9 expression was completely inhibited intriguingly, while expression of septin-2, −6 and −7 were unaffected (Figure-2D). Similarly, western blot analysis of MDA-MB-231, JIMT-1 and MCF-7 cells showed that UR214-9 does not alter the protein expression levels of septin-2,6 and −9 family of proteins (Figure-2E) even though their filamental structures are overwhelmingly disrupted. Septin catastrophe phenomenon in cancer cells was further validated using SKOV-3 ovarian cancer cells that upon treatment with UR214-9 (1μM, 48hours) showed complete disruption of septin-2 filaments wherein septin-2 appears to have relocated to cell surface after drug exposure (Figure-2F). Further examination of septin-6, 7 and −9 structures in drug treated SKOV-3 cells showed reorganization of septin-9 (Figure-2G, lower). Septin-6 was found to be non-needle-like and decreased upon treatment with UR214-9 (Figure-2G, upper). Changes in septin-7 expression were not clear due to non-needle-like and diffused/punctated expression of UR214-9 (Figure-2G, middle).

**Figure 2:**
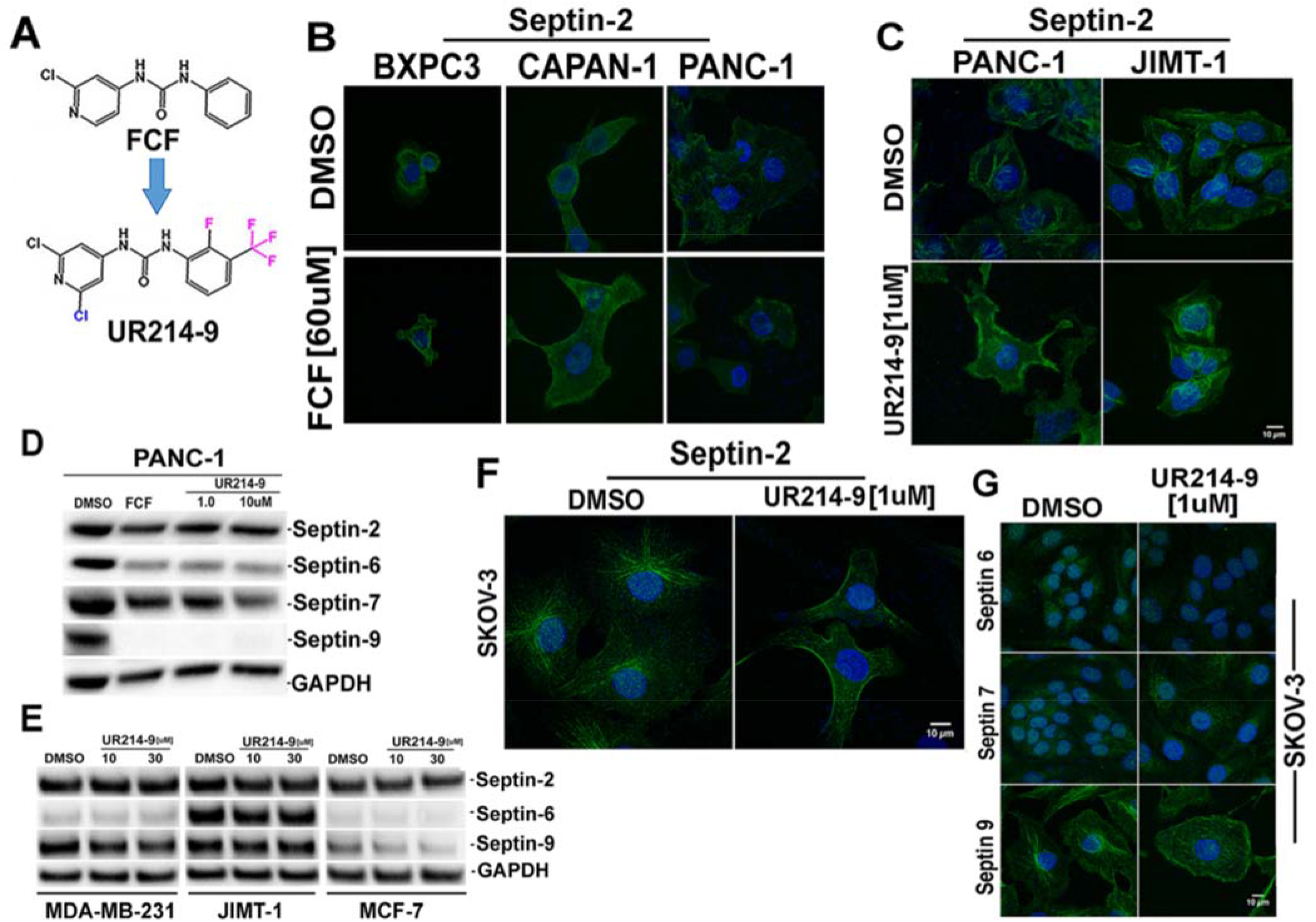
(**A**): Chemical structure of FCF and UR214-9. Structural changes to FCF leading to UR214-9 are shown in blue and pink color. (**B**): BXPC-3, CAPAN-1 and PANC-1 cells were seeded on glass slides and treated with DMSO or FCF(60uM) for 48 hours. The cells were fixed, processed and stained with validated septin-2 antibody (Sigma Aldrich Inc. Catalog number: HPA018481) and source matched secondary antibody (DyLight 488, Vector Laboratories Inc., catalog number: Dl-1488), and confocal images were recorded at 60×2 magnification. (**C**): PANC-1 and JIMT-1 seeded on glass chamber slides cells were treated with DMSO (vehicle) or UR214-9 (1μM) for 48 hrs. Cells were fixed, permeabilized and stained with septin-2 antibody (Sigma Aldrich Inc. Catalog number: HPA018481) and DyLight 488 conjugated secondary antibody (Vector Laboratories Inc., catalog number: Dl-1488), and confocal images were recorded at 60×2 magnification. (**D**): PANC-1 cells were seeded in 100mm3 dishes and treated with DMSO, FCF(60μM), UR214-9 (1.0 and 10.0 μM) for 48 hours. Total cells lysates were immunoblotted and probed with the validated septin-2 (Sigma Aldrich Inc. Catalog number: HPA018481),6 (Sigma Aldrich Inc. Catalog number: HPA005665), 7(Sigma Aldrich Inc. Catalog number: HPA029524) and −9 (Sigma Aldrich Inc. Catalog number: HPA042564) antibodies. (**E**): MDA-MT-231, JIMT-1 and MCF-7 cells seeded in 100mm3 petri dishes were treated with DMSO or UR21409(10 and 30μM) for 48 hours. The total cell lysates were immunoblotted and probed with validated septin-2 (Sigma Aldrich Inc. Catalog number: HPA018481), 6 (Sigma Aldrich Inc. Catalog number: HPA005665) and −9 (Sigma Aldrich Inc. Catalog number: HPA042564) antibodies. (**F-G**): SKOV-3 ovarian cancer cells seeded on glass chamber slides cells were treated with DMSO (vehicle) or UR214-9 (1μM) for 48 hrs. Cells were fixed, permeabilized and stained with validated septin-2 (Sigma Aldrich Inc. Catalog number: HPA018481),6 (Sigma Aldrich Inc. Catalog number: HPA005665), 7(Sigma Aldrich Inc. Catalog number: HPA029524) and −9 (Sigma Aldrich Inc. Catalog number: HPA042564) antibodies followed by DyLight 488 conjugated secondary antibody (Vector Laboratories Inc., catalog number: Dl-1488), and confocal images were recorded at 60×2 magnification.

### UR214-9 causes actin filamental disruption in pancreatic and breast cancer cells

Septins have been previously shown to control the function of actin^22^. Confocal microscopy of UR214-9 treated PANC-1 and JIMT1 cells exhibited actin filament disruption (Figure-3) when treated at a dose of 1μM for 48 hours. Representative structural disarrangement of actin filamental needles, for both PANC-1 and JIMT1 cells, Figure-3. Area of interest are shown in shown in the white boxes.

**Figure 3:**
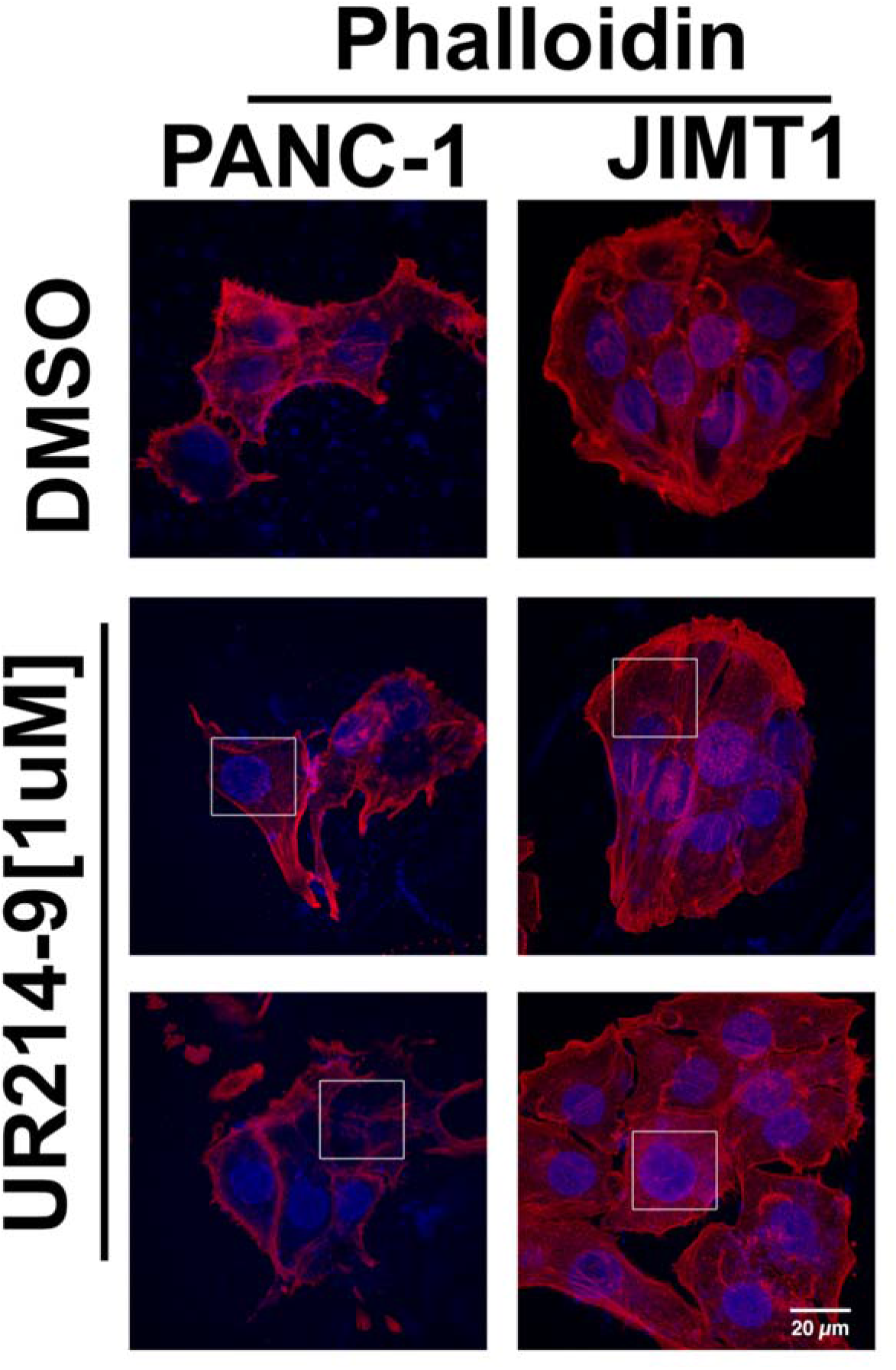
PANC-1 pancreatic cancer cells and JIMT-1 breast cancer cells were treated with vehicle or UR214-9 (1μM) for 48-hour duration, fixed, permeabilized and stained with Phalloidin-TRITC (ECM Biosciences Inc., catalog number: PF7551). Confocal images were recorded at 60×2 magnification. Areas of interest are shown by white boxes.

### *In Silico* docking shows key interactions of UR214-9 with septin-2

Docking experiments to investigate the potential binding mode of UR214-9 and related compounds (including FCF) were performed. UR214-9 and its analogs are smaller in size and similar in structure and symmetry; they consist of a central urea group flanked by two lipophilic substituted aromatic rings. Compounds UR214-8, 9, and 10 are the most active compounds that we have developed; taking this into account, we hypothesized that they could share a similar binding mode (the structure of compounds UR214-8, 9 and 10 are shown in Supplementary Figure 3). Thus, compounds FCF, UR214-8, UR214-9, and UR214-10 were docked into the nucleotide binding site of PDB ID 2QNR, which is the highest quality structure of a septin-2 dimer complex available^23^. Upon visual inspection of the docking poses, two sets of low energy poses (“set A, upper and lower” and “set B, upper and lower”) were identified in which all highly active compounds are able to adopt similar conformations. The two sets are similar to each other in that the three main portions of the molecules – the central urea moiety, the pyridine and the phenyl ring – are in roughly the same area, with the pyridine ring taking the place of the guanine in GDP (Figure 4). In set A, the pyridine nitrogen atom is seen taking the place of the guanine carbonyl oxygen atom, making a hydrogen bond with the backbone of G241 of chain A. ICM scores of set-A were found to be compound UR214-8:-8.85, compound UR214-9: −8.59, compound UR214-10: −10.4 and FCF: −7.21 indicating stronger binding energy of the synthesized analogs than the parent FCF. Set B appears to be identical to a previously reported docking pose for FCF in the same structure template, obtained with the Autodock software^23^. In Figure-4C and D, the identity of amino acid residues interacting with atoms of UR214-9 or its analogs are shown.

**Figure 4:**
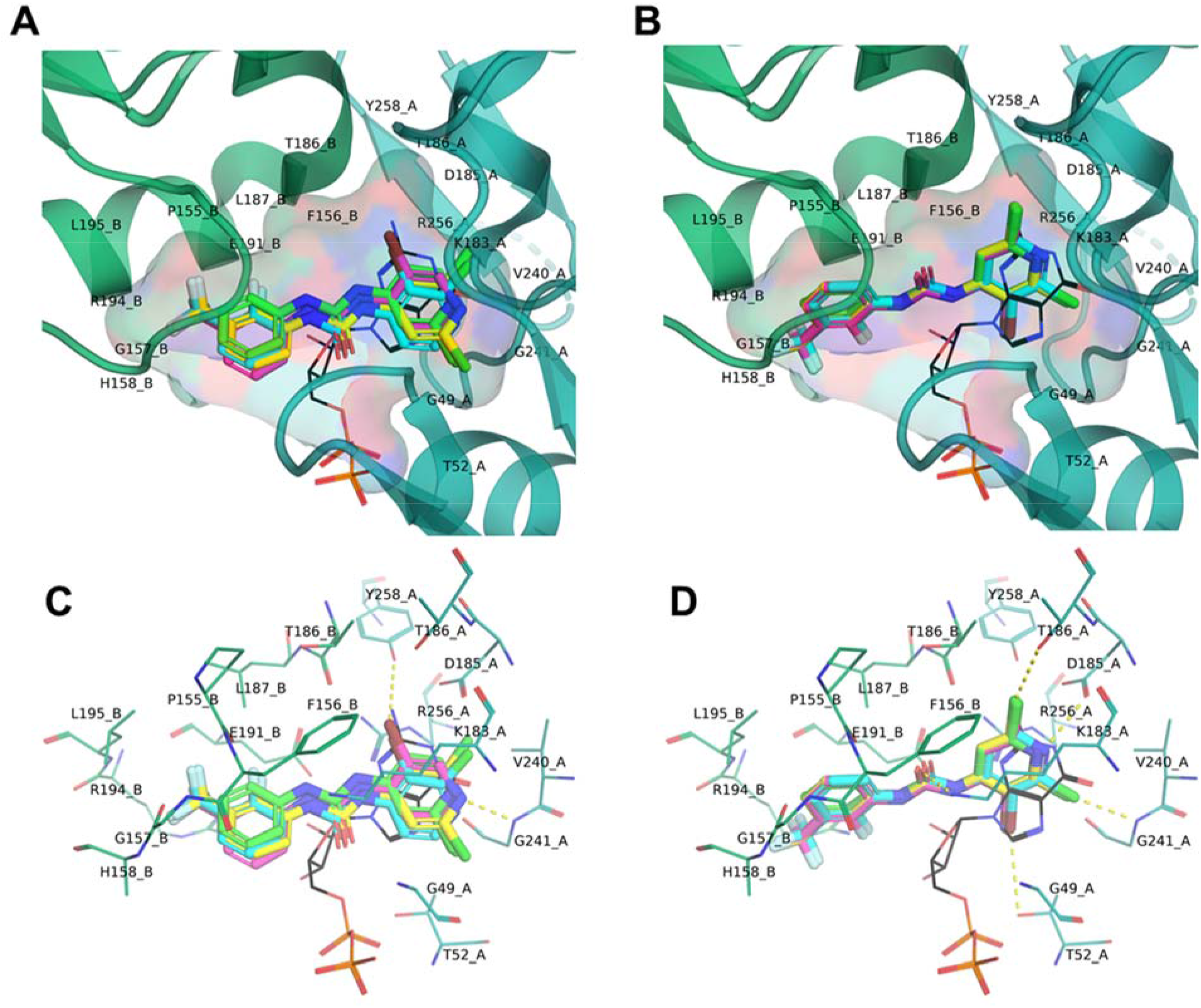
(**A-B**): Septin-2:septin-2 dimer complex was docked with FCF, UR214-8, UR214-9 and UR214-10 using Molsoft’s ICM software package (v. 3.8-7). Compounds were docked into the nucleotide binding site of PDB ID 2QNR, which is the highest quality structure of a septin-2 dimer complex available [33]. Receptor preparation (based on the GDP binding site in chain A) and ligand construction was performed within ICM using standard settings. The compounds were docked with the “dock table” functionality, with a setting for effort of 2.0 and 20 poses per compound. (**C-D**): Amino acid residues interacting with UR214-9 in GDP binding domain are shown.

### UR214-9 impairs cancer cell viability and blocks cell cycle progression

Treatment with UR214-9 reduced the viability of human pancreatic cancer cells (BXPC-3 and PANC-1) cells (Figure 4A) during 72 hours of treatment. PANC-1 and BXPC-3 cells upon treatment with UR214-9 exhibited a large population of non-viable cells based upon staining by the Live-Dead cell kit and by flow cytometry following 72 hours of treatment (Figure 4B and -4C). Given the role of septins in the cell cycle process, we analyzed the effect of UR214-9 on cell cycle progression of PANC-1 and BXPC-3 pancreatic cancer cells at a non-cytotoxic concentration of 100nM. Treatment with UR214-9 at 100nM dose caused minor S-phase arrest in PANC-1 while BXPC-3 cells showed no change in cell cycle distribution at the non-toxic doses (Supplementary Figure-2A). Increasing the dose to 3μM concentration of UR214-9 caused overwhelming arrest in G1 phase (~95% compared to 21%) of BXPC-3 cells, while PANC-1 cells showed complete arrest in sub-G1/G0 phase (data not shown). Similarly, JIMT-1 cells treated with an increased dose (3μM) of UR214-9 exhibited G1 phase arrest and showed largely increased accumulation in G0-phase (Supplementary Figure-2B)

### Analysis of cell cycle protein expression

The spotted antibody array was employed to simultaneous study multiple cell-cycle related proteins expressed in drug treated or naïve PANC-1 cells. Measurement of relative photon counts showed that Cullin-3, glycogen synthase kinase-3 (GSK-3b), p19ARF, 14.3.3.Pan, APC11, APC2, ATM, C-able, CD14Aphsophatase, CDC25C, CDC34, CDC37, CDC47, CDC7, CDH1, CDK1 and CDK-3 were the most expressed and affected proteins in the treated vs naïve PANC-1 considering ≥2.0 fold change as meaningful (Supplementary Figure-3C). β-actin showed the most pronounced expression but expression levels remained unchanged after treatment. On the other hand, Cullin-3 showed most pronounced upregulation in the treated versus naïve PANC-1 cells (Supplementary Figure-3, upper). Cullin-3, a member of the cullin-based ubiquitin ligase family interacts with Hrt1 and BTB domain containing proteins. The resulting complex functions as a Cullin3-based E3 ligase to bring specific substrates to ubiquitinylation and degradation^24^ indicating the role of septins in suppression of cullin-3 mediated ubiquitinylation and subsequent degradation.

### UR214-9 treatment slows the growth of HER2 +xenograft tumors

Septin-2 regulates HER2 expression in gastric cancer cells^25^. HER2 is over-expressed in diverse variety of malignancies^25–27^ and is known to promote tumor development, progression, metastasis and chemoresistance^28^. Septins are shown to protect and stabilize HER2 receptor at the plasma membrane of tumor cells to perpetuate the HER2 orchestrated tumorigenesis^29^. We postulated that targeting septin-2 can potentially emerge as a novel approach to control HER2 orchestrated tumorigenesis. A MTS assay showed that UR214-9 treatment reduced the growth of BXPC-3 and PANC-1 pancreatic cancer dose dependently by 48^th^ hour of drug exposure (Figure-5A). Treatment with UR214-9[3μM] created 29.4% dead cells during 48 hours of drug exposure when the total population of BXPC-3 cells was analyzed by Live-dead kit (Invitrogen Inc). Similarly, PANC-1 cells presented over 38% dead-cell population upon treatment with UR214-9 [3μM]. Next, we determined the impact of UR214-9 treatment on pancreatic cancer xenograft tumor growth *in vivo*. Mice xenografted with HER2+ PANC-1 cells showed significantly delayed growth (p≤0.0001) (Figure-5D). The antitumor efficacy of UR214-9 was further evaluated against HER2 positive xenografts derived from JIMT1 (breast cancer) cells. In addition to increased cell death of JIMT-1 cells upon UR214-9 exposure *in vitro* (Figure-5E), JIMT1 xenograft tumors treated with UR214-9 casted a significant growth control (Figure-5F), based on both tumor volume and weight measurements (Figure-5G).

**Figure-5:**
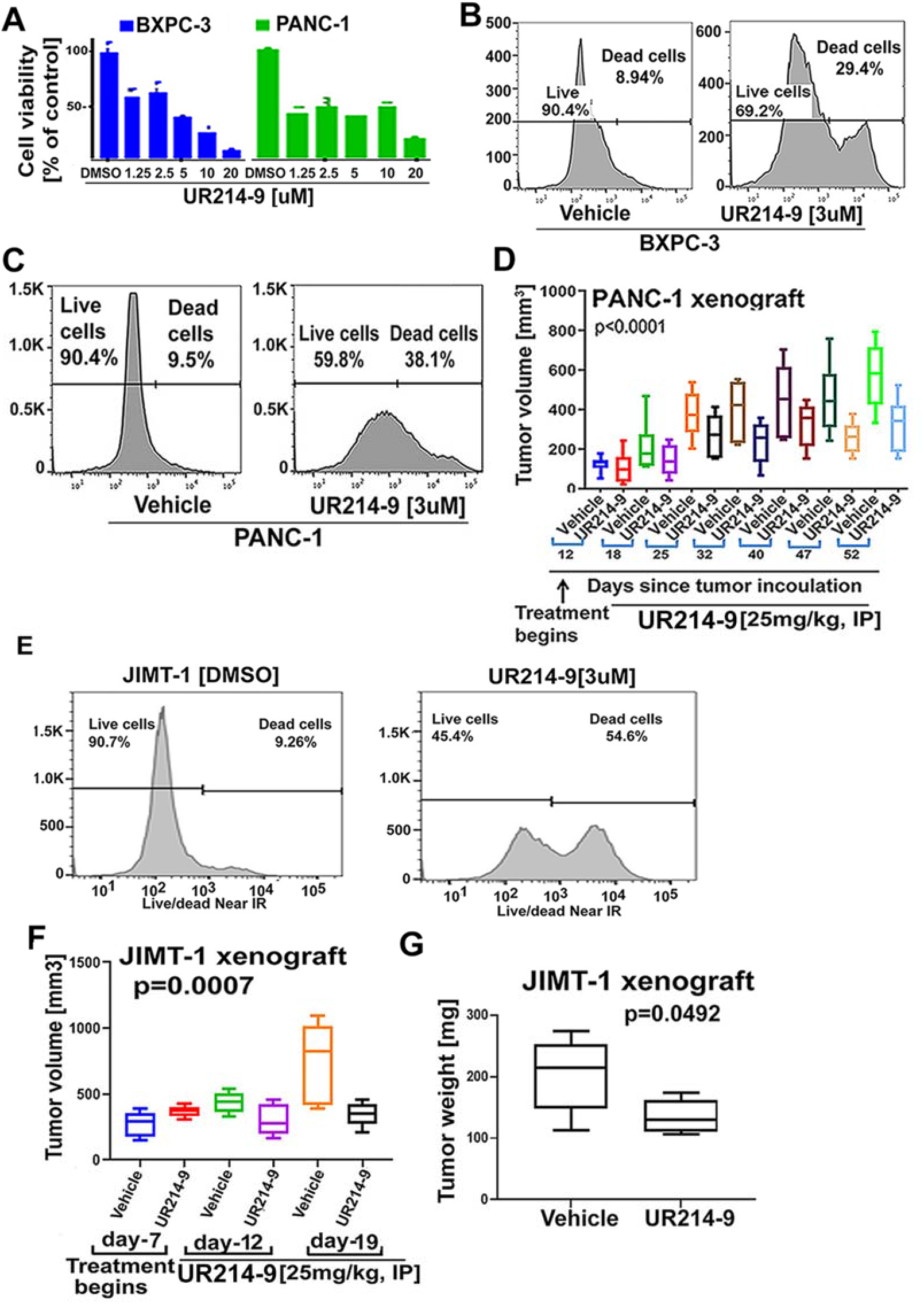
(**A**): Cell viability of PANC-1 and BXPC-3 cells treated with UR214-9 (DMSO, 1.25, 2.5, 5, 10 and 20uM) for 72 hours. The cell viability of the treated groups in comparison with DMSO group was assessed by use of MTS assay (Promega Corp. catalog number: G3580) and absorbance was read at 490nM using BioRad microplate reader. (**B-C**): PANC-1 and BXPC-3 cells were treated with UR214-9(3uM) or DMSO for 48 hours. The cells were stained with Live/dead near IR dye and the live and dead population in vehicle and control group was estimated by flow-cytometry. (**D**): NSG mice (n=10) were inoculated with PANC-1 cells (1 million/animal). Once tumors were palpable, mice were divided into two groups of n=5 each and treated with vehicle or UR214-9 (25mg, M-F, IP, once daily) for 52 days. The tumor sizes were measured periodically. (**E**): JIMT1 cells were treated with UR214-9(3uM) or DMSO for 48 hours. The cells were stained with Live/dead near IR dye and the live and dead population in vehicle and control group was estimated by flow-cytometry. (**F**): NSG mice (n=10) were inoculated with JIMT1 cells (1 million/animal). Once tumors were palpable, mice were divided into two groups of n=5 each and treated with vehicle or UR214-9 (25mg, M-F, IP, once daily) for 28 days. The tumor sizes were measured periodically. (**G**): Tumors from both animals of the control and treatment groups were harvested and weighed on a calibrated balance. Statistical analysis was carried out using GraphPad Prism 8 software. A P value less than 0.05 was considered significant.

### UR214-9 inhibits HER2 expression and blocks phosphorylation of STAT-3

Immunoblot analysis of the total cell lysates of pancreatic cancer cells PANC-1 (HER2+), cells treated with UR214-9 for 72 hours showed a dose-dependent decrease in HER2 expression in PANC-1 (Figure 6A, upper). STAT3 phosphorylation is a down-stream readout of HER2 activation^33^, accordingly UR214-9 treatment also reduced phosphorylated STAT-3 in PANC-1 (Figure 6A, lower) cells. Similarly, UR214-9 treatment reduced HER2 expression in a panel of MDA-MB-231, JIMT-1 and MCF-7 breast cancer cells (Figure-6B) and reduced phosphorylation of STAT-3 in each cell-lines (Figure-6C). We have recently shown that septin-2 is highly overexpressed in ovarian cancer^30^. Similar to JIMT-1 and PANC-1 cell-lines, SKOV-3, a platinum resistant ovarian cancer cell-lines is characterized by HER2 amplification^31–32^. We, therefore employed SKOV-3 cell-line derived xenografts to validate the antitumor efficacy of UR214-9 against HER2 amplified xenograft tumors. To further ascertain the outcome of the combination of UR214-9 with Herceptin, mice were additionally treated with Herceptin alone or in combination with UR214-9. As shown in the Figure-6D, both UR214-9 and Herceptin controlled the growth of tumors. The combination clearly, controlled the tumor growth to a greater degree than both the drugs alone. The real benefit of combination of UR214-9 with Herceptin became apparent when treatments were stopped and tumors were allowed to grow. As shown in the Figure-6D while tumor sizes in UR214-9 and Herceptin group reached the average size in control when the treatments were stopped, the combination maintained greater control over tumor growth (combination p<0.0001**** vs p=0.0004*** and 0.0001*** for vehicle vs UR214-9 and Herceptin). When extracted tumors were weighed, the combination group exhibited presence of smaller tumors, whereas, both the UR214-9 and Herceptin group produced the tumors that matched the average size seen in vehicle group (Figure-6E).

**Figure 6:**
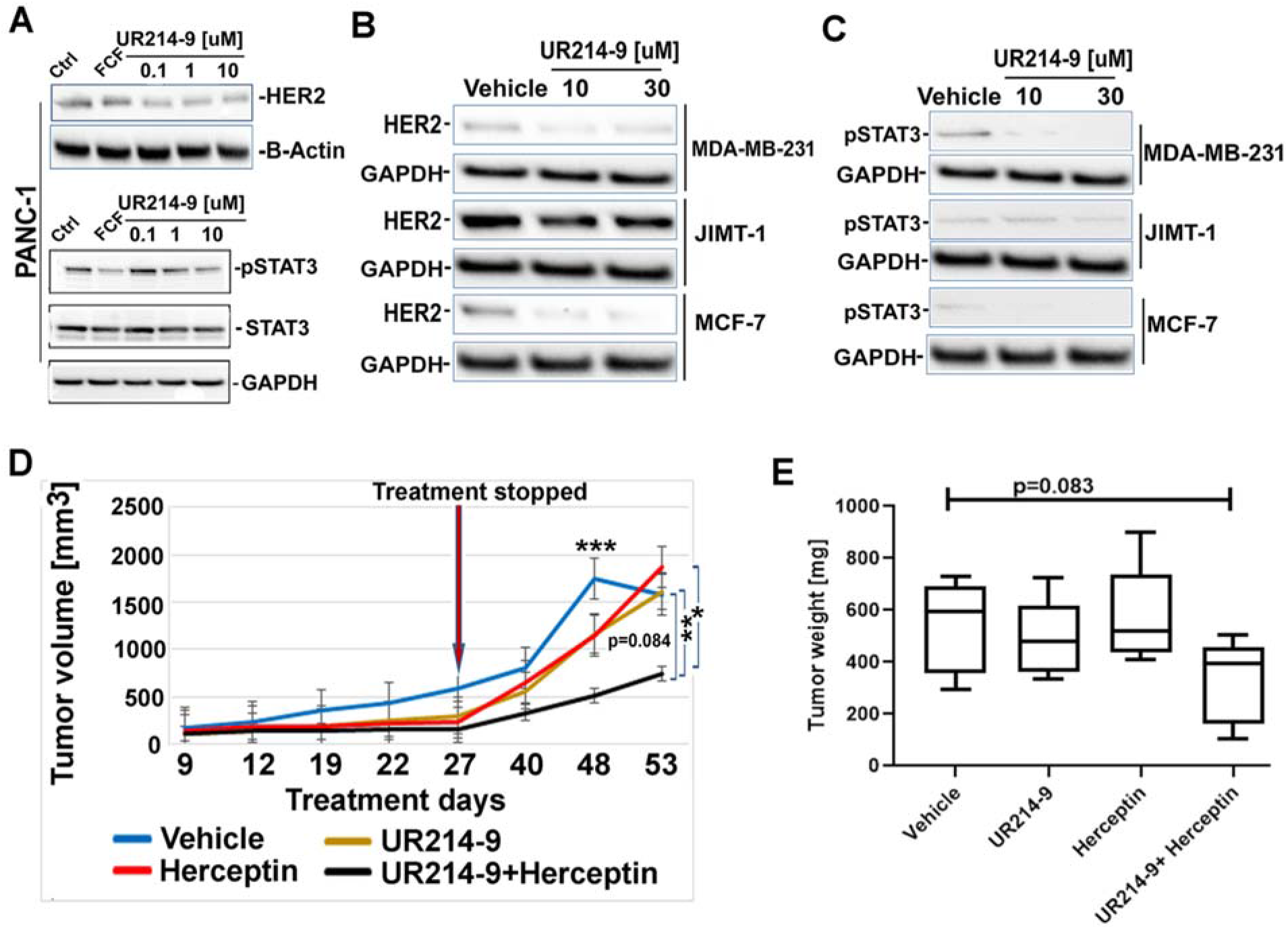
**(A-B)** Adhered BXPC-3, and PANC-1 cells were treated with FCF(60uM) and UR214-9 (DMSO, 0.1, 1.0 and 10uM) for 48 hours. Cells were lysed and immunoblotted with HER2 (Cell Signaling Technology Inc., catalog number: 4290); pSTAT-3 (Cell Signaling Technology Inc., catalog number: 9145p), STAT-3 (Cell Signaling Technology Inc., catalog number: 4904), AKT (Cell Signaling Technology Inc., catalog number: 9272) and GAPDH antibodies (Cell Signaling Technology Inc., catalog number: 2118s). (**C**): MDA-MB-231, JIMT-1 and MCF-7 cells were treated with DMSO or UR214-9 (10 and 30μM) for 24 hours. The cells were lysed, immunoblotted and probed with HER2 (Cell Signaling Technology Inc, catalog number: 4290) and phospho-STAT-3 (Cell Signaling Technology Inc., catalog number: 9145p) antibodies. (D): HER2 expressing SKOV-3 cells (500,000 cells/animals) were implanted in the right flanks of NSG mice subcutaneously. When palpable, mice in groups (n=5 each) were treated with vehicle, UR214-9 (25mg/kg, M-F, I.P.), Herceptin (10mg/kg, M, I.P.). Fourth group was treated with both UR214-9 (25mg/kg, M-F, I.P.) and Herceptin (10mg/kg, M, I.P). Tumor sizes were measured on regular intervals using a digital caliper. Longest length and width were recorded. Tumor volumes was calculated using formula (L*W^2)*0.5 where L represents longest diameter and W stands for width of the tumors measured through a digital caliper. Treatment was stopped on day-27^th^ and tumor sizes were measured on the days indicated. On the day-53^rd^ since inoculation, mice were euthanized and tumors were harvested. Tumor weights were recorded using a calibrated balance. The statistical analyses were performed using Graph Prims 8.1.1.T-test analyses among groups were performed using Graph Prism 8.1.1. version and p<0.05 was considered significant. Day-22: vehicle vs UR214-9: p=0.0035**; vehicle vs Herceptin: p=0.0011**; vehicle vs UR214-9+Herceptin: p=0.0001***; UR214-9 vs UR214-9+Herceptin: p=0.0059**; Herceptin vs UR214-9+Herceptin: p=0.049*. Day-27: vehicle vs UR214-9: p=0.0004***; vehicle vs Herceptin: p=0.0002***; vehicle vs UR214-9+Herceptin: p<0.0001****.

### Whole transcriptome analysis reveals that UR214-9 is target selective

RNA-Seq was performed in the JIMT-1 and Panc-1 cell lines with three treatment groups (10nM Afatinib, 1μM UR214-9, and DMSO) of four replicates each. The samples were sequenced to an average depth of 58 million reads and greater than 90% of the read data for each sample aligned uniquely to the human reference genome (hg38) after adapter and quality trimming. The drug treatments were compared to the control group and differentially expressed genes were determined (adjusted p-value < 0.05). There were 1236 (713 up and 523 down) dysregulated genes between Afatinib treatment and control (Figure 7A and 7C). The ENRICHR webtool was used to determine that the upregulated genes (ALPP, TRIM29, CYP1A1) are associated with extracellular matrix organization and cadherin binding, while the down regulated genes (EGR1, DUSP6, HMGA2, etc.) are associated with purine metabolism and ribosome biogenesis. Conversely, only 11 (7 up and 4 down) genes were called dysregulated between UR214-9 treatment and control (Figure 7B and 7D). In terms of the PANC-1 cell line, there were only two genes (COL13A1 and PRSS22) determined to be significantly differentially expressed upon Afatinib treatment compared to the control group and no differentially expressed genes was called between UR214-9 and the control.

**Figure-7:**
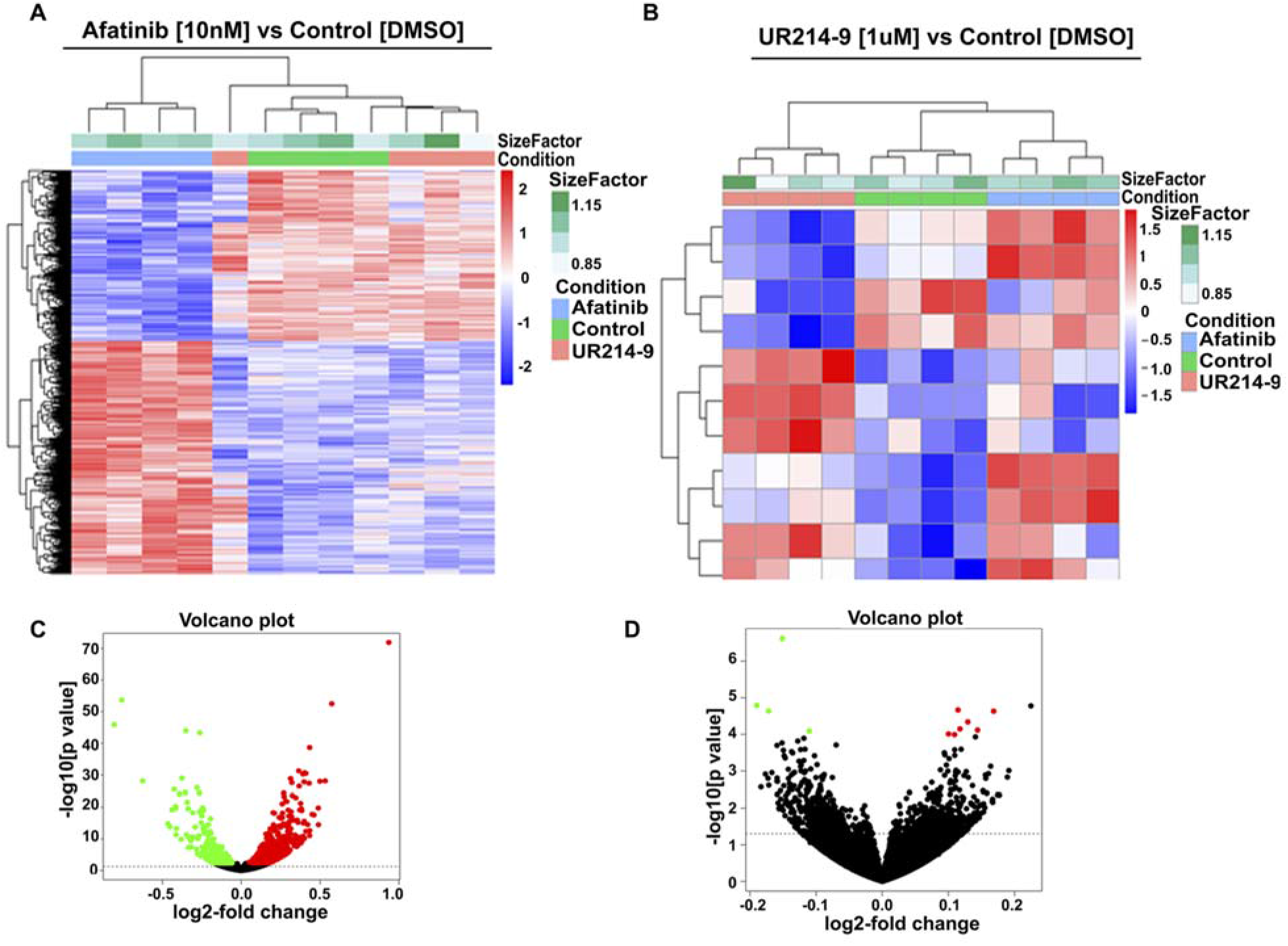
Hierarchically clustered heat map of mRNA expression for 1234 significantly differentially expressed genes (BH adjusted p-value < 0.05) in the JIMT-1 breast cancer cells treated with afatinib compared to control (**A**) and associated volcano plot (**C**). Hierarchically clustered heat map of mRNA expression for 11 significantly differentially expressed genes (BH adjusted p-value < 0.05) in the JIMT-1 breast cancer cells treated with UR214-9 compared to control (**B**) and associated volcano plot (**D**). Heatmap color key represents row scaling of the rLog transformed expression values. The volcano plots have horizontal lines at p-value 0.05 and individual genes/dots are colored red when the adjusted p-value is >= 0.05 and the fold-change is > 0 and green when the fold-change is < 0.

## Methods

### Cell lines, cell culture and reagents

PANC-1, BXPC-3 and CAPAN-1, SKOV-3, MCF7, MDA-MD-231 cells were obtained from ATCC and maintained in DMEM, RPMI-1640 and IMDM supplemented with 10% fetal calf serum penicillin (100 units/mL), and streptomycin (100 μg/mL) at 37°C with 5% CO_2_ in a humidified incubator. JIMT-1 cells were purchased from AddexBio Inc, USA (catalog number: C0006005) and maintained in 10% FBS and antibiotic supplemented DMEM. Septin-2 (catalog number: HPA018481), septin-7 (catalog number: HPA029524), septin-9 (catalog number: HPA029524) antibodies were purchased from Sigma Aldrich Inc. DyLight 488 (catalog number: DI-1488, rabbit, Dylight594 (catalog number: DI-2594, mouse) were purchased from Vector Laboratories Inc. Phalloidin-TRITC was purchased from ECM Biosciences (catalog number: PF7551). HER2 (Cell Signaling Technology, catalog number: 4290); pSTAT-3 (catalog number: 9145p), STAT-3 (catalog number: 4904) and GAPDH antibodies (catalog number: 2118s) were purchased from Cell Signaling Technology Inc. USA and used at manufacturer recommended dilutions.

### Synthesis of FCF analogs

UR214-9 was synthesized by coupling aryl isocyanates with 2,6-dichloro 4-aminopyridines in (0.1:0.1) molar ratio in dry DMF at 65°C overnight under an argon flushed atmosphere. The reaction was monitored using thin-layer chromatography plates with DCM-MeOH or pure ethyl acetate as eluent. Spots were monitored in a UV chamber. The reaction mixture, upon completion of the reaction, was poured into wet ice mixture and triturated and the separated solid was filtered under vacuum. The product was washed with hexane, followed by diethyl ether, and was dried under vacuum. The compounds were characterized by mass spectrometry.

### Molecular docking

Docking experiments to investigate the potential binding mode of 9 and related compounds were performed using Molsoft’s ICM software package (v. 3.8-7). The molecules are rather small and somewhat symmetric (consisting of a central urea group flanked by two lipophilicly substituted aromatic rings). We assumed that since compounds 8, 9, and 10 are the most active ones, that they might share a similar binding mode. Thus, compounds FCF, UR214-8,- 9, and -10 were docked into the nucleotide binding site of PDB ID 2QNR, which is the highest quality structure of a septin-2 dimer complex available to date^23^. Receptor preparation (based on the GDP binding site in chain A) and ligand construction was performed within ICM using standard settings. ICM scores for each compound and their poses were calculated and compared with FCF. The compounds were docked with the “dock table” functionality, with a setting for effort of 2.0 and 20 poses per compound. Upon visual inspection of the docking poses, two sets of low energy poses (“set A” and “set B”) stood out, in which the highly active compounds are able to adopt similar conformations^23^.

### Cell Viability and cell cycle analysis

Cell viability of PANC-1, BXPC-3 and CAPAN-1 pancreatic cancer cells treated with UR214-9 was measured using the Cell Titre96^R^ Aqueous One Solution Cell Proliferation Assay (Promega Corp., catalog number: G3580) following the procedure published earlier. The Live/Dead dye kit (Invitrogen Coro., catalog number: L34975) was used to estimate live and dead cell population in PANC-1 and BXPC-3 pancreatic cancer cells treated with UR214-9 or vehicle. Briefly, cells were treated with vehicle or UR214-9 (3μM) for 72 hours. Cells were harvested by trypsinization, fixed and permeabilized using Fixation-Permeabilization reagent (prepared by diluting the concentrate in the diluent in the ratio 1: 3) (Biogem Inc., diluent: catalog number 92160-00-160 and concentrate catalog number: 2550-00-50) and stained with Live/dead dye for one hour. The cells were centrifuged at 1000 rpm for 5 minutes and pellets were washed and spun down three times with DPBS. The cells were analyzed by a flow cytometer and relative live and dead cell population was calculated by inputing equal number of cells in both vehicle and control group.

For cell cycle analysis, BXPC-3 and PANC-1 and JIMT-1 cells (100,000/well) were seeded overnight in a 6 well dishes and allowed to adhere overnight. Media was replaced with fresh complete medium supplemented with DMSO or UR214-9 (100nM and 3μM) and cells were incubated for 72 hours. The media containing the drugs was removed and cells were washed twice with PBS and trypsinized gently. The cells were collected in 15 mL tubes, complete DMEM media was added to block trypsin and cells were centrifuged. The supernatants were removed and cells were gently treated with 70% cold-EtOH for 30 minutes. The fixed cells were centrifuged and the pellets obtained were collected in flow cytometry tubes and stained with preformulated PI/RNase solution (Cell Signaling Technology, catalog number: 4087s) for 30 minutes. The PI content was analyzed using a flow cytometer. Data was processed using Flowjo software.

### Cell cycle protein expression

Cell Cycle Antibody Array (FullMoon BioSystems Inc, catalogue number: ACC:058), a high-throughput ELISA based antibody array, designed for qualitative/semi-quantitative protein expression profiling was employed to investigate the protein changes after drug treatment. PANC-1 cells were lysed in buffer containing protease and phosphatase inhibitors (Cell Signaling, catalog number: 9803S). Total protein content was quantified by Bradford assay and equal amounts of proteins were analyzed in duplicate with arrays containing 4 to 6 spots for each of 60 probes (ACC058, Cell Cycle Antibody Array; Full Moon Biosystems, Sunnyvale, CA), according to manufacturer’s instructions. After background correction, mean signal intensities were measured using FullMoon Inc’s imaging services. Protein expressions in both the naïve and treatment group was normalized to GAPDH signals.

### Confocal analysis of septin disarrangement

To determine the impact of UR214-9 treatment on Septin-2 structure in cells, PANC-1 or JIMT-1 cells were seeded on glass slides and allowed to adhere overnight. The media was replaced with complete DMEM media supplemented with DMSO or UR214-9 (1μM and 70nM) and cells were incubated for 48 hours. Media was replaced again with new complete medium and fixed with neutral buffered formalin for 15 minutes at 4°C. Media was removed and cells were washed repeatedly with PBST (5× 5mL). The cells were stained with Septin-2 antibody (Sigma Aldrich, catalog number: HPA018481) in PSB overnight at 4°C. Media was removed again and cells were washed with 2×5mL PBST. The cells were stained with fluorescence linked secondary antibody for 1hr under dark. Slides were washed repeatedly in dark for 7×5mL PBST, mounting medium containing DAPI (Vector labs) were applied and covered with glass slide. The slides were stored in dark at 4°C till analysis. Confocal images were obtained and processed essentially as published earlier^34^. Pancreatic tumor microarray (US Biomax, cat no: T142a) were deparaffinized, processed and stained with Septin-2 antibody (Sigma Aldrich, cat number: HPA018481) overnight, washed with PBST and incubated with source matched secondary (FITC) for an hour. Slide was washed in PBST (5×10mL) for five minutes each. DAPI containing mounting media was applied and covered with a glass slide. Confocal images were acquired with Nikon C1si confocal microscope (Nikon Inc. Mellville NY.) using diode lasers 402, 488 and 561. Serial optical sections were obtained with EZ-C1 computer software (Nikon Inc. Mellville, NY). Z series sections were collected at 0.3μm with a 40× PlanApo lens and a scan zoom of 2 or with a 60× Plan Apo objective and a scan zoom of 2, collected every 0.25 μm. Deconvolution measurements were performed with Elements (Nikon Inc. Mellville, NY) computer software. Five cells were outlined and analyzed per field.

### Xenograft studies to evaluate antitumor response of UR214-9

NSG mice were implanted in their left flank with 1 million PANC-1 (HER2+, n=12), JIMT1 (number of animals=10) and SKOV-3 (number of animals=10) cells each in matrigel:media (1:1). Mice were randomized, identified with ear punches and subdivided into vehicle and treatment groups when tumors were found palpable. Both JIMT1 and SKOV-3 formed aggressive tumors within a week and were treated with vehicle or UR214-9 (25mg/kg, IP, seven days a week). PANC-1 formed slow growing tumors and when tumors reached length exceeding 5mm, the treatment was started. A group of SKOV-3 cells were also treated with Herceptin or Herceptin+UR214-9. The vehicle formulation was: 40% Hydroxypropyl-beta-cyclodextrin [Acros Organics] & solutol HS15 (Sigma] in sterile water). 25mg/kg equivalent of UR214-9 (1uL=200ug in DMSO) was dissolved in 600uL PBS+400uL of the vehicle and vortexed to obtain a clear suspension. Tumor burden and animal weight was measured manually by digital calipers on weekly or biweekly routine. Tumor volume was calculated using the formula ½(L × W^2) where L is a longest diameter and W is the widest width. Statistical difference between the vehicle and treatment groups was analyzed by GraphPrism-8 software using one way annova. P<0.05 was considered significant. Mice after the treatment period were euthanized and tumors were resected, weighed and frozen in liquid nitrogen. A portion of the tumors from the control and treatment groups were fixed in neutral buffered formaldehyde and paraffin embedded. 5uM thickness tissues slides were prepared for histochemistry.

### mRNA Sequencing

The total RNA concentration was determined with the NanopDrop 1000 spectrophotometer (NanoDrop, Wilmington, DE) and RNA quality assessed with the Agilent Bioanalyzer (Agilent, Santa Clara, CA)^35^. The TruSeq Stranded mRNA Sample Preparation Kit (Illumina, San Diego, CA) was used for next generation sequencing library construction per manufacturer’s protocols. Briefly, mRNA was purified from 200ng total RNA with oligo-dT magnetic beads and fragmented. First-strand cDNA synthesis was performed with random hexamer priming followed by second-strand cDNA synthesis using dUTP incorporation for strand marking. End repair and 3’ adenylation was then performed on the double stranded cDNA. Illumina adaptors were ligated to both ends of the cDNA, purified by gel electrophoresis and amplified with PCR primers specific to the adaptor sequences to generate cDNA amplicons of approximately 200-500bp in size. The amplified libraries were hybridized to the Illumina single end flow cell and amplified using the cBot (Illumina, San Diego, CA). Single end reads of 75nt were generated for each sample using Illumina’s NextSeq550^36^.

### Whole transcriptome data analysis

Raw reads generated from the NovaSeq6000 sequencer were demultiplexed using bcl2fastq version 2.19.0. Quality filtering and adapter removal are performed using Trimmomatic-0.36 with the following parameters: “TRAILING:13 LEADING:13 ILLUMINACLIP:adapters.fasta:2:30:10 SLIDINGWINDOW:4:20 MINLEN:35” Processed/cleaned reads were then mapped to the *Homo sapiens* reference sequence (GRCh38, hg38) with STAR-2.6.0c given the following parameters: “--twopassMode Basic --runMode alignReads --genomeDir ${GENOME} --readFilesIn ${SAMPLE} --outSAMtype BAM SortedByCoordinate --outSAMstrandField intronMotif --outFilterIntronMotifs RemoveNoncanonical”. The subread-1.6.1 package (featureCounts) was used to derive gene counts given the following parameters: “-s 2 -t exon -g gene_name”. Differential expression analysis and data normalization was performed using DESeq2-1.16.1 with an adjusted p-value threshold of 0.05 within an R-3.4.1 environment. Heatmaps were created using the pheatmap R package^37–39^.

### Data acquisition and statistical analysis

The prognostic assessment of septin-2, −7 and −9 in the panel of different cancers was conducted using Human Protein Atlas tools. Alternatively, R2 genome.org tools were employed to determine the impact of septins enrichment on the survival prospects. P values less than 0.05 were considered significant. The relative tumor sizes in the naïve vs treat groups were calculated using GraphPrism8 using one-way annova settings. P values less than 0.05 were considered significant.

## Discussion

Data continue to emerge on the association of septins with malignancies, making identification of septin-targeted therapies crucial to block the aberrant septin functions in cancer cells. Considering FCF that essentially strengthens septin-2^40^ as the starting point, we have developed UR214-9, a small molecule which dismantles septin-2 and −9 filamental assembly in cancer cells without killing the cells or altering the septins protein levels in the cells. To our best knowledge, this is the first description of a pharmacologic approach to disrupt septin’s structure in cells. We considered that disrupting oligomeric septin filamental structures via UR214-9 treatment will be the key to impact cytokinesis and control the proliferation of cancer cells. Not only did the disruption of septin filaments by UR214-9 reduce the proliferation of pancreatic cancer cells (and of breast, ovarian endometrial, lung and kidney cancers; some data not shown) in vitro, xenografted tumors of breast, ovarian, pancreatic and lung malignancies (data not shown) treated with UR214-9 also showed reduction in tumor growth. Interestingly, the combination with Herceptin led to stronger control over HER2 positive SKOV-3 xenograft’s growth (Figure-6D).

Enhancement in antitumor effects of Herceptin via co-treatment with UR214-9 in HER2 positive ovarian cancer xenograft model is stemming likely from the association of septin-2 with HER2. Septin-2 is shown to maintain HER2 signaling in cancer cells^27^. Septins protect and stabilize HER2 receptor at the plasma membrane of tumor cells to perpetuate the HER2 orchestrated oncogenic signaling and tumorigenesis^27^. It is anticipated that targeting septins can improve the survival rate of HER2 positive breast, pancreatic and other malignancies such as ovarian and lung. HER2 overexpression leads to aggressive breast malignancy and poor patient survival^41^. Current repertoire of therapies for HER2+ malignancies are inadequate. More than 60% of HER2+ breast cancer patients do not respond to trastuzumab treatment and resistance to the treatment develops rapidly in virtually all patients^42^. Further, the inability of trastuzumab to penetrate solid breast tumors to block secreted (truncated) forms of HER2, that promote resistance and metastasis, limits its usefulness in providing a complete and lasting control over HER2 orchestrated breast tumor growth^43^. Similarly, treating or preventing brain metastases in patients with HER2+ breast cancer is challenging^44^, particularly in the post-trastuzumab phase of treatment. About two-thirds of patients develop brain metastases despite control or response of their extracranial disease to trastuzumab^45^. Because trastuzumab does not penetrate the central nervous system, the brain may serve as a sanctuary site^46^. A blood-brain barrier (BBB) penetrant drug would be required to better control brain metastases in patients with HER2 +positive cancers. UR214-9 carries the structural attributes of small polar surface area signatures (calculated for UR214-9=53.49 vs <90 required to cross BBB) that would facilitate penetration through the BBB. UR214-9, therefore, may improve outcomes of patients with brain metastases from HER2+ cancers.

Signaling associations of septins are not fully understood. To determine signaling association of septins and perturbations that UR214-9 treatment mounts, we conducted global rna-seq analyses of breast and pancreatic cancer cells treated with UR214-9 and, as a comparator, afatinib, a HER2 targeted therapy. As shown in Figure-7, afatinib treatment clearly had the most impact on the transcriptional profile of PANC-1 cells, while treatment with DMSO and UR214-9 did not have much effect on the transcriptome. The lack of differentially expressed genes between the UR214-9 treatment and control suggests that the mode of action of UR214-9 is non-transcriptional, and treatment with UR214-9 does not appear to elicit a gross transcriptional response.

Taken together, this study demonstrates that aberrant septins expression indicates poor prognoses among patients with cancer. UR214-9 is the first prototype of a small molecule that can induce septin-2 and −9 filamental catastrophe, a pharmacologic and cytoskeletal response of the cells not described before, to control cancer cell proliferation and tumor growth. Moreover, an important pharmacologic feature of UR214-9 is the benefits of limited off-target engagements. As shown in Figure-7, compared to afatinib, an EGFR targeted therapy that affected gene expression of over 1200 genes in JIMT-1 breast cancer cells, UR214-9 treatment even at 100-fold higher dose affected less than 20 genes significantly. Although UR214-9 is a close structural analog of FCF, UR214-9 differs significantly from FCF pharmacologically. While FCF is shown to strengthen septin-2, UR214-9 dismantles septin-2 and septin-9 filamental assembly. ICM scores calculated through molecular docking indicated greater binding affinity of UR214-9 with septin-2:septin-2 dimer complex than FCF. To the best of our knowledge, other than FCF, which is clinically unfit due to the weak pharmacologic effects, off-target effects and functions associated with strengthening the septin-2 filaments, UR214-9 is the only septin modulator described so far that can dismantle septin’s structural arrangement in nano molar concentrations (70nM-1uM). Given the preliminary antitumor response in breast, pancreatic, ovarian and lung cancer xenograft models (data not shown) and its therapeutic capabilities to significantly enhance the response of Herceptin in HER2 expressing xenograft tumors it is apparent that dismantling septins is an effective and clinically promising approach to prevent tumor growth, although doses, delivery formulations and frequencies of administrations have to be optimized, and a synergistic or at least an additive combinational agent has to be identified to achieve fuller control over the tumor growth. Based on the promising outcome in combination with Herceptin, our laboratory is currently evaluating the outcome of combination of UR214-9 with paclitaxel and Herceptin in breast cancer models to increase the clinical utilities of UR214-9.

---

**Supplementary Figure-1:**
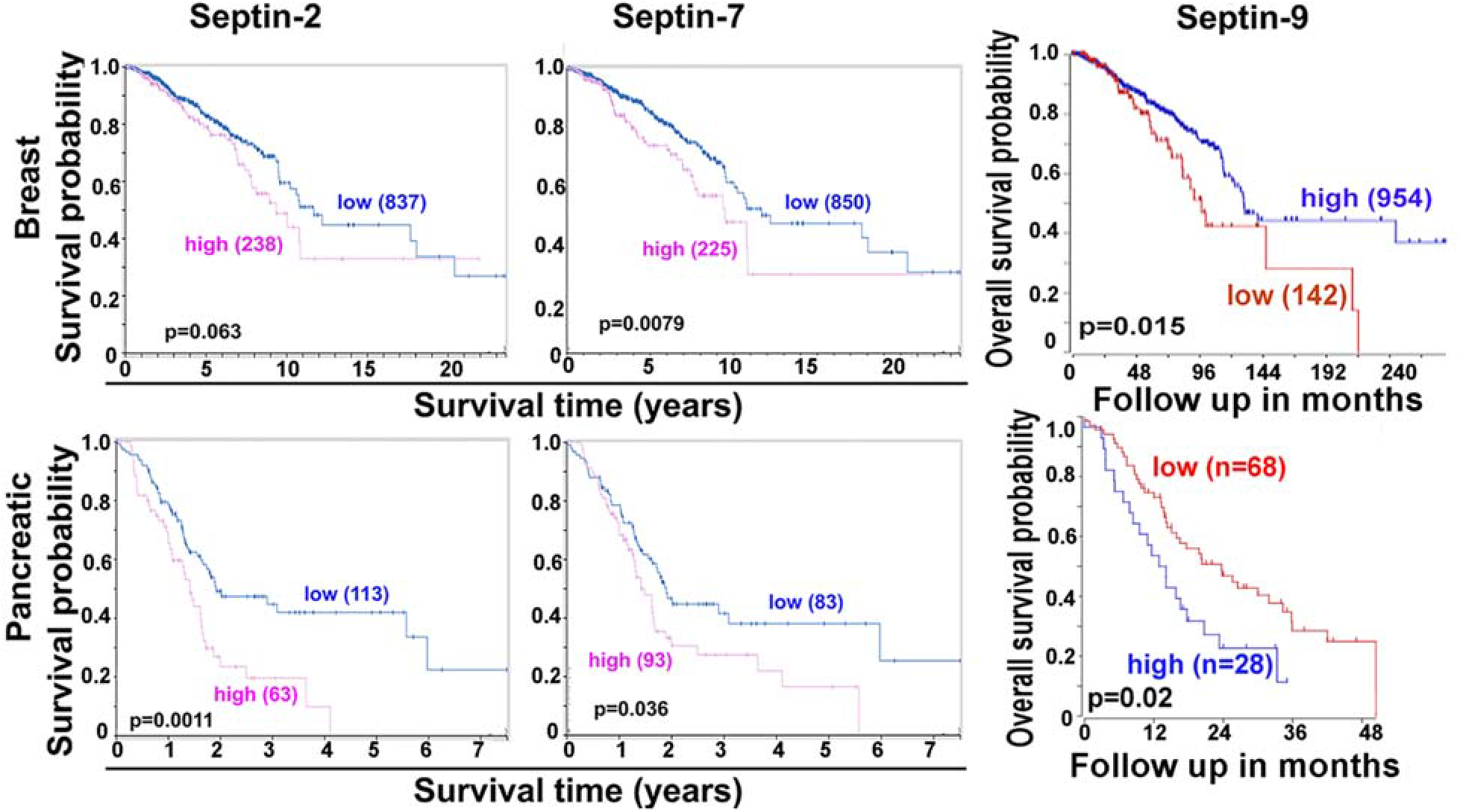
Survival probabilities of patients diagnosed with breast and pancreatic, was correlated with septin-2 (left), septin-7 (middle) and septin-9 (right) gene expression using the data and tools available at the Human Protein Atlas (https://www.proteinatlas.org/) or R2-Genomics Analysis and Visualization Platform (https://hgserver1.amc.nl/cgi-bin/r2/main.cgi). P values less than 0.05 were considered significant.

**Supplementary Figure-2:**
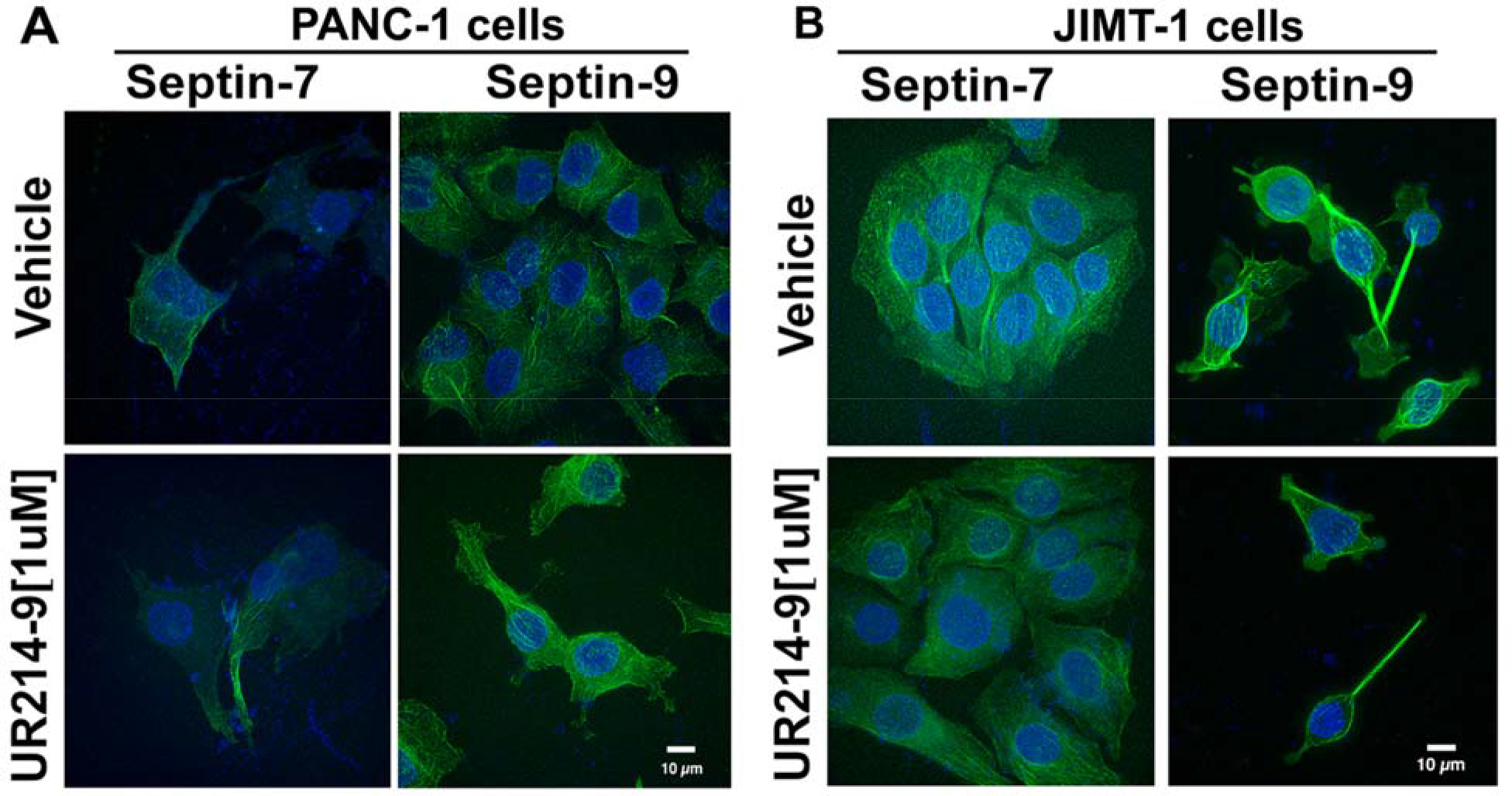
PANC-1 and JIMT1 cells were treated with DMSO or UR214-9 (1μM) for 48 hrs. Cells were fixed, permealized and stained with septin-7 and −9 antibodies (Sigma Aldrich Inc., catalog number: HPA029524, HPA042564) and DyLight 488 conjugated secondary antibody (Vector Laboratories Inc., catalog number: Dl-1488), and confocal images were recorded at 60×2 magnification.

**Supplementary Figure-3:**
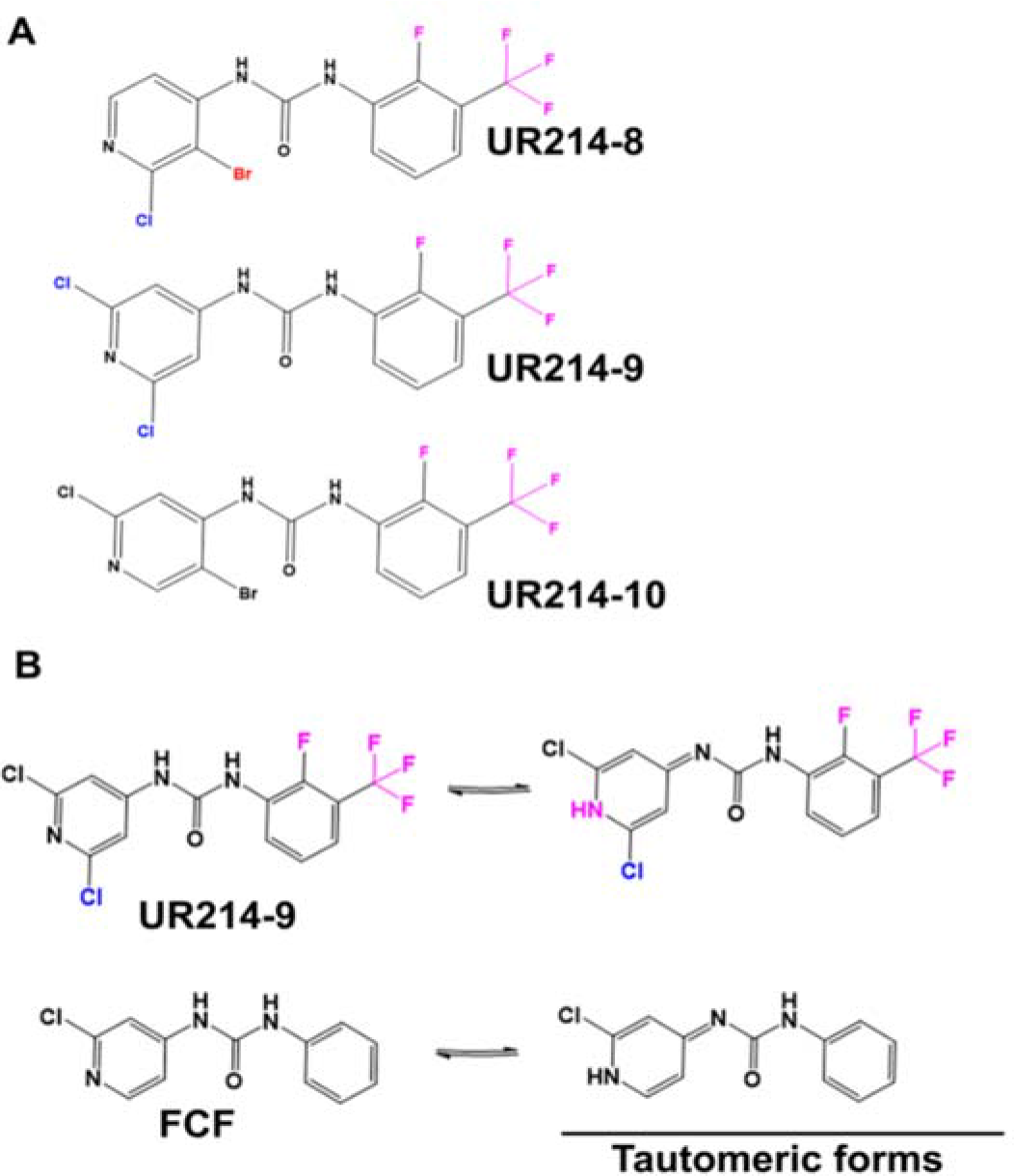
(**A**): Chemical structures of UR214-9 analogs employed in molecular docking. (**B**): Possible tautomeric forms of UR214-9 and FCF.

**Supplementary Figure-4:**
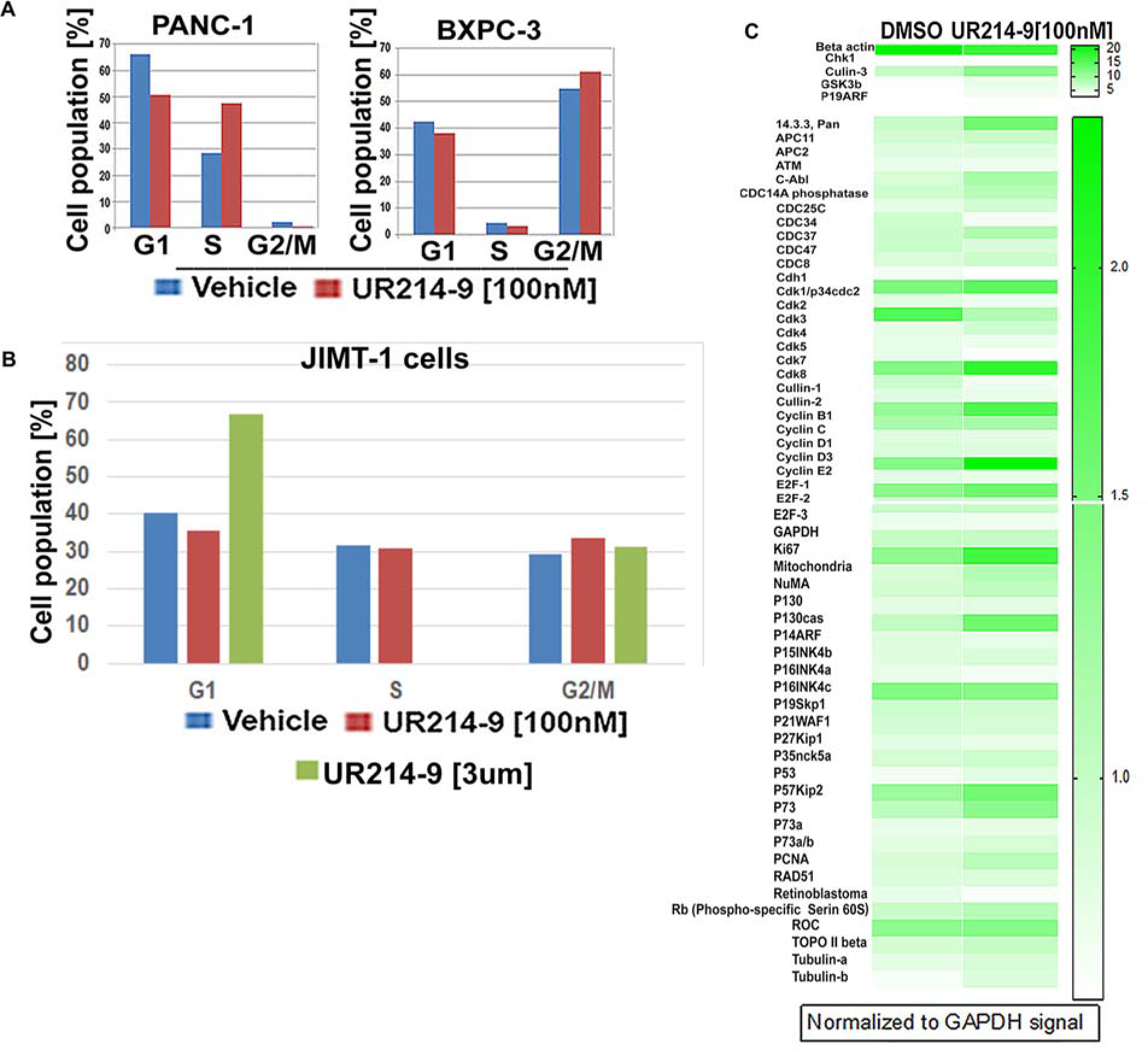
**(A):** PANC-1 and BXPC-3 cells were treated with sub-cytotoxic concentrations of UR214-9 (100nM for 48 hours). Cells were fixed, permeabilized and stained with pre-formulated PI-RNase solution (Cell Signaling Technology, catalog number 4087s). DNA content was measured using flow-cytometry and cell cycle distribution was analyzed by flowjo or FCF express software. (**B**): JIMT1 cells were treated with [(DMSO, UR214-9 (0, 100nM-3μM)] for 48 hours, fixed, permeabilized and stained with pre-formulated PI-RNase solution (Cell Signaling Technology, catalog number 4087s). DNA content was measured using flow-cytometry and cell cycle distribution was analyzed by flowjo software. (**C**): PANC-1 cells were treated with DMSO or UR214-9 (100nM) or 3μM for 48 hours. The cells were lysed and total protein was isolated using the buffer available in FullMoon Bioscience Cell Cycle Antibody array kit (catalog number: ASC058). The proteins isolated from the vehicle and treatment groups were applied on the antibody array, and processed and developed per the manufacturer’s instructions. The photons were read using the array Image Quantification and Analysis services of FullMoon BioSciences (Catalog number SDA01-ACC058) and normalized to GAPDH. Cullin-3, GSK3b and p19ARF followed by Pan 14.3.3 were the most upregulated proteins in the treatment groups. Heatmap shows fold-changes in protein expressions.

## Availability of data and UR214-9

The complete set of in vitro and in vivo results, rna-seq and western blot data are available from the corresponding author upon request. Similarly, reasonable quantities of UR214-9 will be freely made available for research and studies upon request.

## Author contributions

RS conceived the idea, designed study, synthesized compounds including UR214-9 and conducted animal experiments in team with LL. LL ran xenograft animal studies, recorded tumor size measurements and animal weights without involvements of RS. LR and JK participated in animal studies. CL conducted the docking studies. AJ, NK, PS, RP, AA and KKK conducted western blot, immunoprecipitation, cell viability and other supplementary studies. VH performed confocal microscopic studies. TC conducted antibody array, cell cycle and live-dead cell assay experiments. RNA-seq data was generated and analyzed by the team of CB, JRM, EZ and JA. RS assembled the manuscript. RT, MTM, DL, RGM and SG reviewed the data and edited the manuscript. Every author read and edited the manuscript, and approved the present version of manuscript for submission.

## Acknowledgements

KKK and RS are grateful to UR Ventures of University of Rochester for a pilot grant award. RS gratefully acknowledges Mae Goode Foundation award to partially support these studies.

## Conflict of Interest

KKK, RBRT, RGM and RS are listed as the co-inventors on a provisional US patent application US62/894,424 covering UR214-9 and its analogs for the treatment of malignancies and other disease states orchestrated by septins.

